# Pharmacological manipulation of olfactory bulb granule cell excitability modulates beta oscillations: Testing a model

**DOI:** 10.1101/234625

**Authors:** Boleslaw L. Osinski, Alex Kim, Wenxi Xiao, Nisarg Mehta, Leslie M. Kay

## Abstract

The mammalian olfactory bulb (OB) generates gamma (40 – 100 Hz) and beta (15 – 30 Hz) oscillations of the local field potential (LFP). Gamma oscillations arise at the peak of inhalation supported by dendrodendritic interactions between glutamatergic mitral cells (MCs) and GABAergic granule cells (GCs). Beta oscillations occur in response to odorants in learning or odor sensitization paradigms, but their generation mechanism and function are still poorly understood. When centrifugal inputs to the OB are blocked, beta oscillations disappear, but gamma oscillations persist. Centrifugal input targets primarily GABAergic interneurons in the GC layer (GCL) and regulates GC excitability, which suggests a causal link between beta oscillations and GC excitability. Previous modeling work from our laboratory predicted that convergence of excitatory/inhibitory inputs onto MCs and centrifugal inputs onto GCs can increase GC excitability sufficiently to drive beta oscillations primarily through voltage dependent calcium channel (VDCC) mediated GABA release, independently of NMDA channels. We test this model by examining the influence of NMDA and muscarinic acetylcholine receptors on GC excitability and beta oscillations. Intrabulbar scopolamine (muscarinic antagonist) infusion decreased or completely suppressed odor-evoked beta in response to a strong stimulus, but increased beta power in response to a weak stimulus, as predicted by our model. Piriform cortex (PC) beta power was unchanged. Oxotremorine (muscarinic agonist) tended to suppress all oscillations, probably from over-inhibition. APV, an NMDA receptor antagonist, suppressed gamma oscillations selectively (in OB and PC), lending support to the model’s prediction that beta oscillations can be supported by VDCC mediated currents.

**New and Noteworthy:**

- Olfactory bulb beta oscillations rely on granule cell excitability.
- Reducing granule cell excitability with scopolamine reduces high volatility-induced beta power but increases low volatility-induced beta power.
- Piriform cortex beta oscillations maintain power when olfactory bulb beta power is low, and the system maintains beta band coherence.

## Introduction

In mammalian olfactory systems, several local field potential (LFP) frequency bands have been identified, but thus far only gamma oscillations (40-110 Hz) have been explained from the mechanistic to the functional level. Beta oscillations (15-30 Hz) have previously been shown to arise in response to associative odor discrimination learning and odor sensitization. These oscillations overwhelm the olfactory bulb (OB) and pyriform cortex (PC) LFP during odor sampling, and yet they remain mechanistically and functionally enigmatic. When centrifugal input to the bulb is lesioned, beta oscillations are extinguished, but gamma oscillations persist (Neville and Haberly, 2003; Martin et al., 2006). Beta oscillations also increase in power with the onset of learning in operant tasks (Martin et al., 2004, 2007; Frederick et al., 2016). They are more likely to be involved in mediating the contextual meaning of an odor or in carrying out the action associated with that meaning than in the objective representation of an odor during early sensory processing. We have shown that the timing of these oscillations is related to late odor sampling in operant odor discrimination tasks, emerging in a fast (<100 msec) transition from gamma-dominated activity after 2-4 sniffs of the odor (Frederick et al., 2016). Interestingly, beta oscillations can also be elicited in waking rats without any specific association after repeated exposure to high volatility odorants (Lowry and Kay, 2007). This suggests that there may be multiple functional routes to these oscillations.

In a previous report we introduced a model that addressed the role of granule cell (GC) excitability in generating beta oscillations (Osinski and Kay, 2016). This model was inspired by three lines of evidence: (1) Beta oscillations cannot be generated without intact centrifugal input to the OB (Neville and Haberly, 2003; Martin et al., 2006); (2) centrifugal input largely targets the GC layer and synapses onto GCs are perisomatic (Figure 1A; Mouret et al. 2009, but see Illig, 2011); (3) GCs can undergo at least two types of long lasting afterdepolarizations, one dependent on GC muscarinic receptor activation (Castillo et al., 1999; Pressler et al., 2007) and the other on an appropriate convergence of mitral cell (feedforward) and cortical (feedback) inputs onto GCs (Egger et al., 2005). Another model also predicted that OB beta oscillations can only occur when GCs are in high states of excitability and relied on spiking GCs (David et al., 2015). Some studies have shown that GCs spike rarely and can release GABA in a graded fashion at the dendrodendritic reciprocal synapse (Schoppa et al., 1998; Cang and Isaacson, 2003). We therefore interpreted excitability not as a tendency to spike but rather a tendency to drive graded inhibition (model summarized in Figure 1).

**Figure 1.**
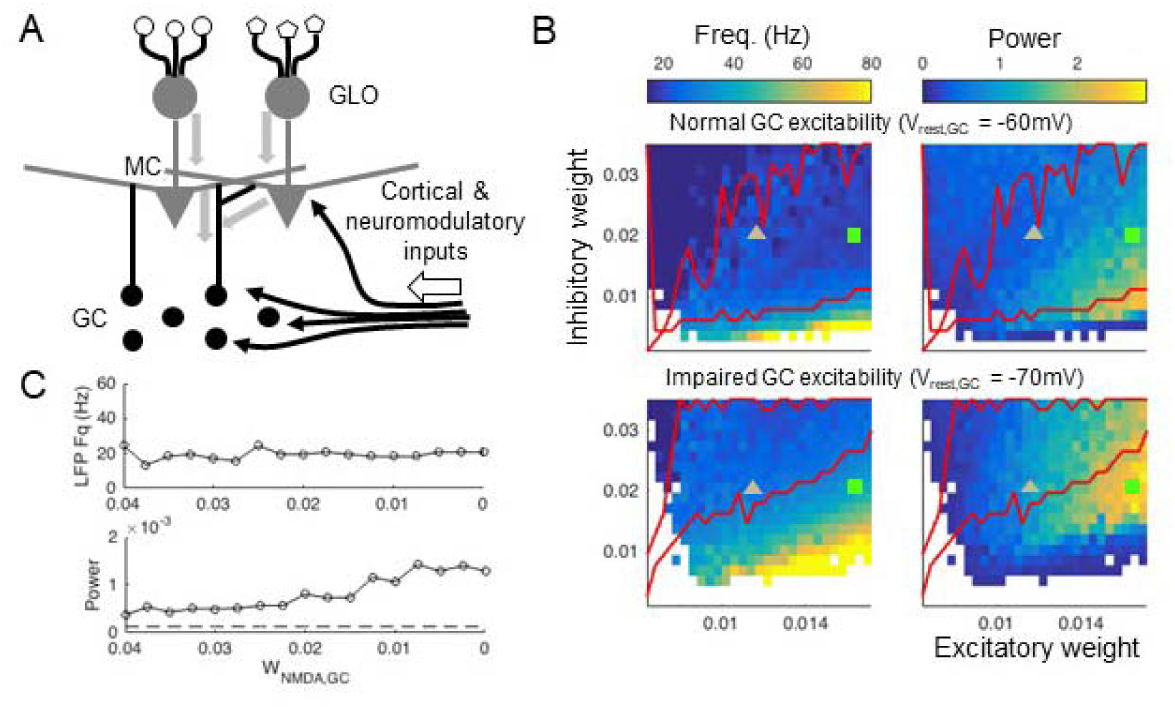
Modeling predictions. **A:** Schematic of olfactory bulb circuitry. Glomeruli (GLO) innervated by unique receptors receive sensory inputs which are conveyed to mitral cells (MCs). Sensory (thick gray arrows) and cortical/neuromodulatory feedback (thin black arrows) both converge onto granule cells (GCs). **B:** Modeling results (Osinski and Kay, 2016) showing odor-induced LFP power and frequency landscapes under normal (top) and impaired (bottom) GC excitability conditions. The y-axis is inhibitory weight, which represents the strength of inhibition from GCs to MCs, and the x-axis shows excitatory weight, which represents the strength of sensory input to MCs. The boundary of the beta regime (20 – 30 Hz) is drawn in red. White regions had power less than or equal to the noise floor, which was defined as the maximum power of the residual oscillation due to common inputs to MCs when inhibition is removed. The MC stimulation due to strong (high volatility) and weak (low volatility) odorant stimuli at constant inhibitory weight are marked by a green square and gray triangle, respectively. In the normal condition (top) the evoked beta frequency is fairly stable with respect to sensory input strength and both low (gray triangle) and high (green square) volatility odorants induce beta oscillations. In the impaired condition (bottom) the GC resting potential (V_rest,GC_ in the model) remains at −70 mV, and the evoked beta frequency is less stable over the same range of inputs. In this condition, the high volatility odorant (green square) generates high power low gamma oscillations, while the low volatility odorant (gray triangle) induces higher power beta oscillations that it did in the normal condition. **C:** (Top) Beta oscillation frequency remains stable as the strength of the NMDA current from GC to MC (W_NMDA,GC_ in the model) is gradually reduced to 0, because they are sustained by the N-type voltage gated channel current. (Bottom) In this particular simulation, the LFP power increases as NMDA is reduced because the system is over-inhibited for high values of W_NMDA,GC_. The dashed line indicates power of the noise floor due to common excitation without inhibition. (Figures adapted and edited from Osinski and Kay (2016), with permission.)

The model predicted that GC excitability influences the stability of OB LFP oscillation frequency. Beta oscillations were generated over a wide range of parameters when GCs were in a high state of excitability, providing sustained graded inhibition onto MCs. Decreasing GC excitability destabilized the LFP frequency, such that stronger stimuli would drive oscillations out of the beta regime, while weaker stimuli could actually induce higher power beta. The bi-directional effect on beta oscillation power was a consequence of shifting the balance of excitation and inhibition onto MCs. This is illustrated in Figure 1B, which shows the range of LFP frequencies generated by the simulated MC-GC dendrodendritic network as a function of sensory input to MCs and strength of GC-MC inhibition in normal and impaired GC excitability conditions. Two points illustrate the predicted effects of low volatility (weak odor, low excitatory weight) and high volatility (strong odor, high excitatory weight) odorants on the LFP under these two conditions. Under normal conditions, a high volatility odorant (green square) induces beta oscillations with relatively high power, and a low volatility odorant (gray triangle) induces lower power beta oscillations (Figure 1,B top). Under impaired GC excitability conditions, the strong odorant induces high power low gamma frequency oscillations instead of beta oscillations (Figure 1,B bottom). Conversely, the weak odorant induces higher power beta oscillations than it did under normal GC excitability. Thus, the model predicts that for strong odors, a drug that lowers GC excitability should attenuate the power of beta oscillations, but for weak odors the same drug should enhance the evoked beta oscillations.

Our model also predicted which channels and currents maintain beta vs. gamma oscillations. Beta oscillations could be sustained in the model by graded inhibitory currents, mediated primarily by voltage-dependent calcium channels (VDCCs), even when NMDA currents were blocked (Figure 1C). Gamma oscillations, however, could not be maintained in the model without intact NMDARs, which matched results showing that APV, an NMDAR antagonist, blocks gamma oscillations in the OB of waking mice (Lepousez and Lledo, 2013).

We tested the predictions of our model experimentally by infusing drugs into the OB that modulate GC excitability in several ways. Muscarinic receptors are found in high density in the EPL, where GCs synapse onto MC lateral dendrites and at moderate densities in the GC layer (Fonseca et al., 1991; Lein et al., 2007). Muscarinic receptor agonists can both inhibit or excite GCs by differential activation of M1 or M2 receptors (Castillo et al., 1999; Mandairon et al., 2006; Smith and Araneda, 2010; Li and Cleland, 2013) and can influence their excitability by modulating afterdepolarization (Nickell and Shipley, 1993; Pressler et al., 2007). When we infused scopolamine, a muscarinic antagonist, into the ventromedial OB prior to an odor sampling session, beta oscillations were decreased in response to a high volatility odorant and increased in response to a low volatility odorant.

Furthermore, we found that infusion of APV, an NMDAR antagonist, suppressed gamma but not beta oscillations. Both of these results align well with the model’s predictions. Injection of oxotremorine, a muscarinic agonist produced more variable results, but in some rats we observed a tendency for OB gamma and beta to be suppressed, possibly due to over-inhibition. Although beta oscillations require intact connections between OB and PC (Neville and Haberly, 2003; Martin et al., 2006), we found that PC beta oscillations show a degree of independence in power, and coherence between the OB and PC is relatively unaffected by decreasing beta in the OB. Together, these results confirm our model predictions and reveal a more nuanced picture of the generation of beta oscillations in the olfactory system than was previously understood.

## Methods

Subjects were 8 adult male Sprague-Dawley rats (350 – 450 g; purchased from Envigo (Harlan)), maintained in the colony room on a 14 −10 h light/dark schedule (lights on at 08:00 CST). Two rats were used for pilot studies to determine drug dosages. Six rats were used for the data as reported. Rats were housed singly after electrode implantation and had access to unlimited food and water for the course of the experiments. All animal procedures were done with approval and oversight by the University of Chicago Animal Care and Use Committee with strict adherence to AAALAC standards.

### Electrode implants

Before each surgery, rats were given a subcutaneous injection of ketamine cocktail (35 mg/kg ketamine, 5 mg/kg xylazine, and 0.75 mg/kg acepromazine). Anesthesia was maintained by checking for reflexes every 15 min and administering intraperitoneal injections of ketamine. Bipolar stainless steel formvar insulated electrodes (100 μηι wire; ~1 mm vertical tip separation) were placed in the left anterior OB (8.9 mm anterior to bregma, 1.5 mm lateral, average depth 3 mm), left aPC (0.5 mm anterior to bregma, 3.0 mm lateral, 15° angle, average depth 8 mm), and right ventromedial OB (8.5 mm anterior to bregma, 1.5 mm lateral, average depth 4 mm) as shown in Figure 2A. A cannula guide (Plastics One C315, 26 gauge, 5 mm guide with 0.75 mm internal injection cannula projection) with two stainless steel electrodes attached on either side of the shaft was implanted in the left ventromedial OB (8.2 mm anterior to bregma, 1.5 mm lateral, average depth 2.5 mm). Bipolar electrodes were vertically positioned across the ventral mitral cell layer in the OB and across the layer 2/3 pyramidal cell layer in the aPC by orienting the electrode perpendicular to the cell layer and lowering it until the LFP was reversed across the two leads of the bipolar electrode. The cannula was inserted into the GCL by observing a reduction in the amplitude of the LFP after lowering past the dorsal mitral cell layer. Internal cannula depths determined *postmortem* are shown in Figure 2D. Reference and ground wires attached to stainless steel screws were secured to the skull over the left cerebellum and right occipital lobe respectively (see REF & GND Fig. 2A). Additional screws were used for securing the headstage to the skull. Connector pins for each lead were inserted into a round plastic receptacle (Ginder Scientific, Ottawa, Canada), and the assembly was embedded in dental acrylic. Rats were allowed to recover for 2 weeks after surgery before beginning the experimental protocol.

**Figure 2.**
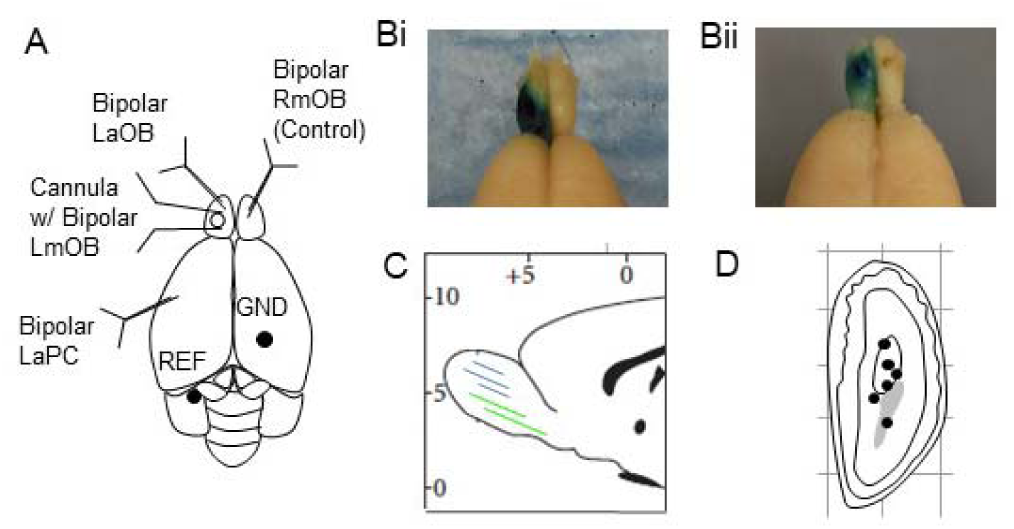
Electrode placement and histology. **A:** Schematic showing electrode and cannula placements. **B:** Two representative examples showing spread of methylene blue dye (injected just before perfusion) localized to the left OB. **C:** Schematic sagittal slice showing approximate anterior-posterior spread of drug (blue lines) as determined by spread of methylene blue dye. (The two green lines correspond to the two rats with large contralateral scopolamine effects whose means are colored green in Figure 6C.) **D:** Schematic coronal section of LmOB showing approximate internal cannula tip depths determined via Prussian Blue reaction.

### Verification of electrode placements and drug spread

After experiments were complete, rats were deeply anesthetized with a euthanizing dose of Urethane (1g/kg), and methylene blue dye was injected into the cannula at the same rate and volume used for drugs during experiments (4 μL, at 1 μL/min). Dye could not be injected into one rat due to cannula blockage. Current was passed between each electrode and the ground screw, thus depositing iron residue at the tips of the stainless steel wires. The brain was fixed via intracardial perfusion, and electrode tips were marked using the Prussian Blue reaction. Prior to sectioning, brains were extracted from the skull, the electrode array was removed, and the brains were sunk in 30% sucrose (0.1 M phosphate buffered formal saline) for two days and then flash frozen in isopentane cooled to −40deg. OB cannula placements were confirmed by visual examination of coronal slices showing electrode tracks and the blue stain marking the electrode tips (Fig. 2D). The spread of methylene blue dye across rats is shown in in Fig. 2B,C.

### Drugs and dosages

We used three drugs in this study: scopolamine (Sigma Aldrich, MW 339.81g/mol), oxotremorine (Sigma Aldrich, MW 322.19 g/mol), and APV (Sigma Aldrich, 197.13 g/mol). We ran pilot experiments on 2 rats (not including the 6 rats used for the study) to determine the appropriate dose of these drugs. For our initial choice of Scopolamine dose we followed (Mandairon et al., 2006), who used a low concentration of 7 mM and a high concentration of 38 mM. In our pilot experiments, we found no effect of low concentration on beta power and small effects at the higher dose. We found that a slight increase to 50 mM produced much stronger effects. In this study we refer to 38mM as the low concentration and 50mM as the high concentration. We did not use the 7 mM dose, because we wanted to limit the number of drug conditions planned for each subject. For APV we tried to follow Leposez & Lledo (2013) who infused 0.25 mM and 1 mM for high and low concentrations. However, both of these doses caused seizures in our rats. We found that 100 μM was a safe dose that still produced significant effects. For oxotremorine, we followed Bendahmane et al. (2016) and Smith et al. (2015) who used 10 μM and 30 μM doses. No significant effects were seen at 10 μM, but 30 μM produced strong effects (though they were not always consistent, see Results). Higher concentrations tended to induce seizures.

Initially, we intended to use low and high doses of each drug, but because oxotremorine and APV produced seizures at higher doses and showed little effect at lower doses in our pilot study, and the need to limit the number of infusions for each subject, we only used low doses of oxotremorine and APV for this study. There were no seizures induced by scopolamine in our pilot rats, even at concentrations higher than those used in the present study.

### Experimental design

After full recovery from surgery, rats were habituated to odors for one day before the start of experiments to eliminate novelty effects. Drugs were freshly mixed immediately before each experiment. At the start of the experiment, rats were plugged into a Multichannel Systems wireless transmitter headstage (W16-HS) and then placed in a clean polycarbonate cage with fresh bedding. Each experiment started with 5 minutes of free behavior recording to obtain a baseline LFP used to normalize the odor evoked gamma and beta oscillations. Afterwards, 4 μL of drug or saline was infused into the left OB at a rate of 1 μL/min. If the cannula was clogged, then 2 μL of saline was carefully injected by hand to clear the blockage. In some cases the cannula became permanently clogged (possibly due to scar tissue and glial cell buildup) and no further sessions could be run, resulting in partial data for those subjects. A control session with saline injection was performed after every two drug sessions to track any systematic changes in the LFP from day to day. The order of drug sessions was balanced across rats.

Drug effects often started immediately after or even during the infusion (Fig. 3C), so we began odor presentations immediately after infusion. Two odorants were used in this study, Ethyl-2-methylbutyrate (EMB, Sigma Aldrich) and Geraniol (GER, Sigma Aldrich). EMB has a high volatility (theoretical vapor pressure-VP- at 25° C of 7.86 mm Hg) and elicits high amplitude beta oscillations, while GER has a low volatility (VP 0.0133 mm Hg) and elicits smaller beta oscillations (Lowry and Kay, 2007). Odors were presented in blocks of 24 trials with ~20 s between trials (total 48 trials). These odor blocks spanned the two phases of an apparent biphasic scopolamine effect (see Figs 3 & 4). For each trial, an odor soaked cotton swab was held under the rat’s nose for two or three sniffs (as judged by the presenter). In two of the rats, we also used an interleaved odor presentation design, where EMB and GER were alternated on each presentation for a total of 48 trials, producing 12 trials of a given odor for each half of the experiment. Interleaving the odor presentations did not change the main results of the experiment. The drug and odor order were balanced across all 6 rats. The timeline of the odor block presentation experiment is depicted in Figure 4B.

**Figure 3.**
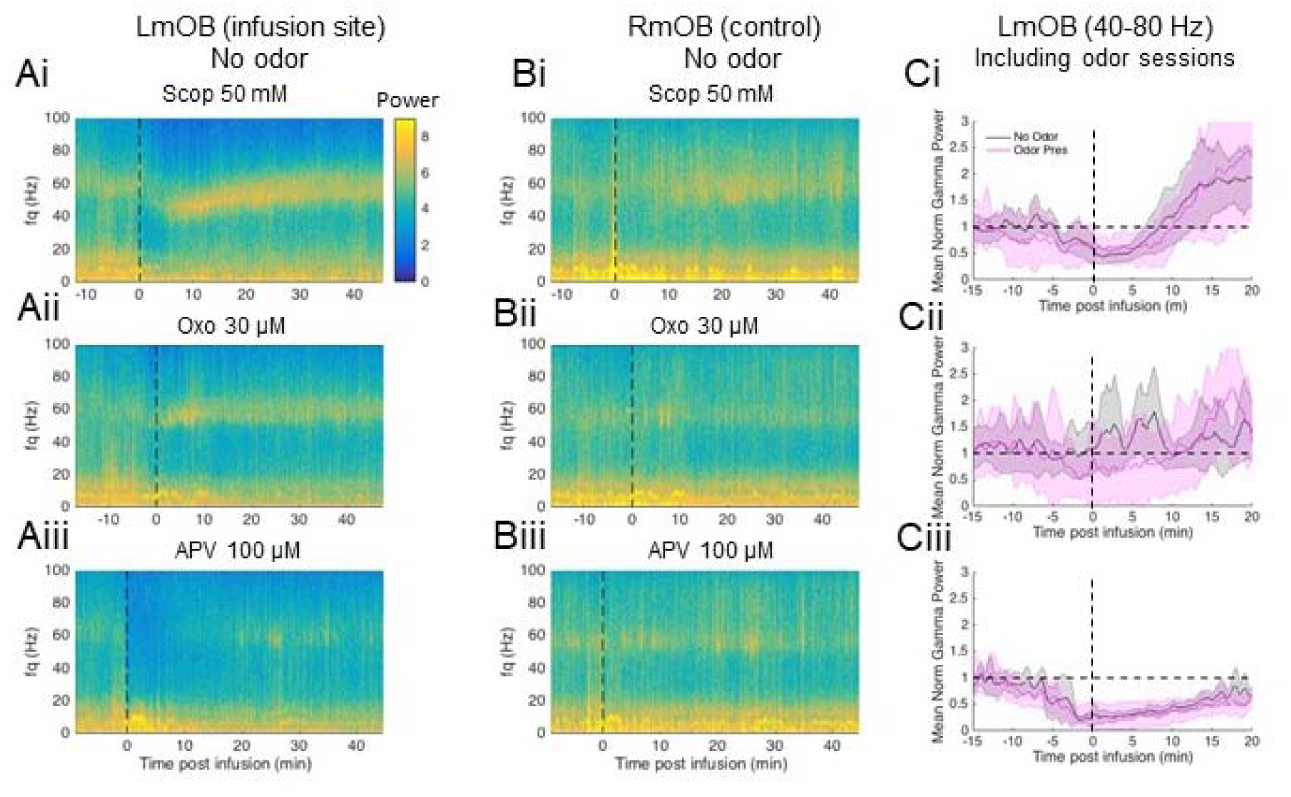
Temporal effects of drugs on OB LFP power. **A:** Representative spectrograms of LmOB LFP after infusion of scopolamine (Ai), oxotremorine (Aii), and APV (Aiii) into the LmOB without any odor presentations. Vertical dashed lines indicate end of infusion (time to complete the infusion varied slightly from session to session). **B:** Spectrograms of contralateral (RmOB) LFP activity corresponding to the sessions in A. **C:** Mean gamma (40 – 80 Hz) power averaged over no odor (gray) and odor presentation (pink) sessions aligned to end of infusion time (vertical dashed lines indicate end of infusion). Power was normalized by the average baseline power (indicated by horizontal dashed line). Scopolamine and APV had fairly consistent effects across subjects, but oxotremorine effects were highly variable.

**Figure 4.**
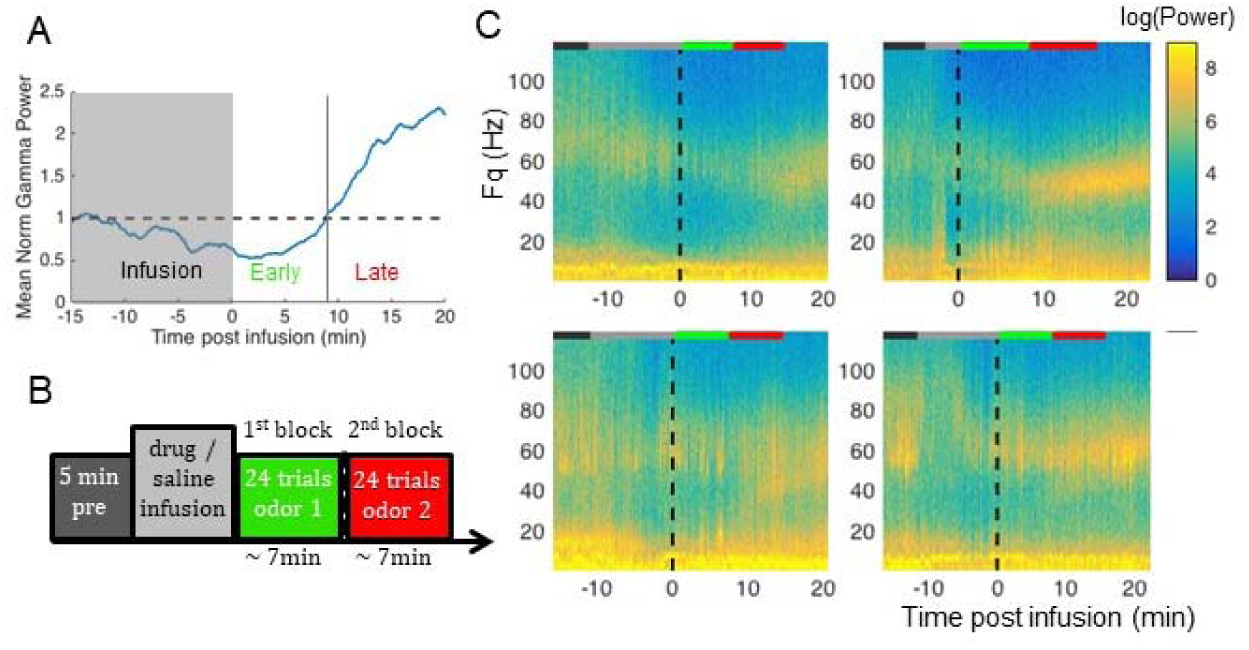
Determining average duration of early phase scopolamine effect in LmOB. **A:** Mean normalized power in the LmOB gamma band (40-80Hz) from all scopolamine sessions aligned by the end of infusion time-points. The infusion period (gray block) differed in duration from session to session (see methods). Dashed horizontal line is the baseline gamma power before infusion (power was normalized by baseline, so baseline is at 1). We interpret the early phase of scopolamine as the time post infusion it takes for mean gamma power to return to baseline (marked by vertical line), which is about 7-8min **B:** Odor presentation experimental timeline. The 1^st^ block overlaps with the early phase of scopolamine effect, the 2 ^nd^ block with the late phase. **C:** Example spectrograms of scopolamine (50 mM) sessions in 4 rats. Vertical black dashed lines indicate end of infusion. Colored lines at the top of each plot indicate the duration of each step in the experiment using same color scheme as the timeline in B. In each of these plots, the 1^st^ (green) and 2^nd^ (red) odor blocks bracket the transition between the two phases of the scopolamine effect. Odor evoked beta and gamma oscillations can be seen in some of the plots. In these plots, the 2^nd^ odor block starts just before the onset of the rebound gamma (scopolamine effect phase 2).

### Electrophysiology

All data were recorded wirelessly using Multichannel Systems 32-channel basic wireless recording system, using a 16-channel headstage (W16-HS) with a digital sampling rate of 2 kHz. Each lead was recorded with reference to a skull screw above the left cerebellum (see REF in Fig. 2A). A ground screw was placed over the right posterior cortex. Approximate odor onset times were recorded with a 5V TTL pulse triggered by the experimenter pressing a button just before the rat took its first sniff. Brain signals and TTL pulses were recorded with MC Rack software (http://www.multichannelsystems.com/downloads/software).

### Spectral analysis

For each rat, we assessed the quality of signals from the two leads from each brain area. We chose the lead with the cleanest signals and most prominent beta oscillations from each pair and used this lead for analysis across the entire set of experiments. All analysis was performed in MATLAB^®^ R2015b. We filtered out movement artifact (MATLAB ‘designfilt’ function using ‘lowpassfir’ with pass 4Hz, stop 8Hz, then applied with the filtfilt function) and subtracted it from the original data. We found that finite impulse response filters produced less prominent edge effects than infinite impulse response. We then extracted the odor presentation trials from the continuous LFP trace in 4-second windows starting from the button press at the start of each trial. We calculated power spectra for each trial using the multitaper method implemented in the Chronux version 2.11 toolbox for MATLAB (Bokil et al., 2010), with a time-half-bandwidth of 2 and 3 tapers over a frequency range of 1 Hz to 100 Hz. This method multiplies each LFP trace with a series of tapers (Slepian sequences) and then averages them, which has the effect reducing spurious noise contributions. The Slepian sequences also possess desirable spectral concentration properties, which produce a more accurate measure of the underlying power spectrum (Bokil et al., 2010). We divided the LFP power spectra into three bands: beta (15 – 30 Hz), low gamma (40 – 60 Hz), and high gamma (60 – 100 Hz). We discarded the first 4 trials of each 24 trial odor block, because beta often only comes on after the 3^rd^ or 4^th^ odor presentation, consistent with findings in Lowry and Kay (2007). We then averaged the power in each frequency band across the remaining 20 trials, and report the peak beta band power from the averaged power spectra.

To calculate coherence between OB and PC signals we used the coherencyc function in the Chronux toolbox. This function applies a multitaper coherence calculation to the entire 4s odor period for each trial and then averages across trials. We used 9 tapers with a time-half bandwidth of 5. We then applied Fisher’s Z transform of the coherence (Kay and Freeman, 1998; Kay and Beshel, 2010), defined as tanh^−1^(coherence), to distribute the values from zero to infinity instead of zero to one.

### Double normalization of power spectra

The LFP amplitudes differed across subjects and days, primarily due to differences in electrode placement and condition across subjects, electrode drift within subjects, and effects of repeated infusions. Normalizing the LFP power during odor presentations by baseline power alone was not sufficient to put the power values of different rats on the same scale for meaningful statistical comparisons, because different rats may show different degrees of odor-evoked oscillatory power increase even under normal conditions. These differences are often due to differences in electrode placement or quality. Therefore, we also compare the relative odor-evoked power changes under drug vs. saline conditions. Thus, we normalized each power spectrum first by the power in the 5 min baseline before the infusion and then by the power during odor presentations in the most recent saline session. The power in the 5 min baseline period was determined by dividing it into 4 s long non-overlapping windows and averaging the power of all these windows. Using the same length windows as for odor presentation ensured that the baseline spectrum had the same frequency resolution as the power of the odor presentation trials. Windows with significant movement artifacts were discarded. The saline sessions were also normalized by their baselines before being used as normalizations for the drug sessions.

### Statistical methods

5-way unbalanced ANOVA was conducted for scopolamine and oxotremorine sessions separately using the MATLAB anovan function. Factors were subject, saline/drug, odor identity, odor block, and frequency band. For APV sessions we conducted a 4-way ANOVA, because odor blocks were combined for analysis of these sessions. An unbalanced design was used because some rats had missing sessions, either due to the cannula becoming permanently blocked or a drug causing seizures even at lower doses (these data were excluded). In addition, in some cases there were different numbers of repetitions (uniform odor blocks had rep = 20, interleaved odor blocks had rep = 10, after excluding the first 4 trials in each block).

When ANOVAs were significant, we performed post-hoc t-tests between each saline and drug pair using the mean for each rat. There were 2 (odors) × 2 (odor blocks) × 3 (frequency ranges) = 12 comparisons for each drug, giving us a Bonferroni correction factor of 12, and setting the significance threshold at p < 0.05/12 ≈ 0.004. (For APV, we found no differences across odor blocks, so the trials were treated as a single block, with the number of comparisons reduced to 6, p < 0.008. See Results- *Effects of drugs on baseline LFP*.) Throughout the paper we present the data in violin plots (using MATLAB File Exchange function violin.m), which show a smoothed probability density distribution (using a Gaussian kernel) of the normalized power in beta, low gamma, and high gamma frequency bands averaged across all rats.

## Results

We designed the experiments to test the hypothesis that manipulating GC excitability would affect the amplitude of beta oscillations as predicted by our model (Fig. 1). Two drugs, scopolamine and oxotremorine, tested the effects of GC modulation by the muscarinic ACh receptor. A third drug, APV, tested our model’s prediction that blocking NMDA receptors would decrease gamma but not beta oscillations.

### Effects of drugs on baseline LFP

We timed odorant presentation to occur within the duration of the drug effects. To measure the duration of the drug effects, we recorded LFPs from 3 rats for 45 min after infusion without any odor presentations. Representative effects on the left medial OB (where the drug was infused) and right medial OB (a control) of a single rat are shown in Fig. 3A,B (end of infusion marked by vertical dashed line). To visualize the average effect we aligned the LFPs of each session to the end of infusion time, and averaged the 40 – 80 Hz gamma band power across subjects (Fig. 3C, gray traces). All three drugs typically began to take effect during infusion. APV (100 μM) had a highly consistent effect in every session, nearly abolishing gamma for over 20 minutes post infusion (Fig. 3Ciii). The muscarinic drugs, however, had more complex effects.

Scopolamine infusion produced a consistent biphasic effect on the LFP, starting with a strong suppression of LFP power across gamma and beta frequencies, followed by a “rebound” boost in low gamma power, which slowly dissipated as gamma crept up to the baseline gamma frequency over about 20 min (Figures 3 Ai and 3 Ci). We measured the duration of the early phase as the time that the average broadband gamma (40 – 80 Hz) power took to return to pre-infusion baseline gamma power levels, which we found to be ~7-8 mins after the end of infusion (Fig. 4A). Because of this biphasic effect we designed the odor presentations to fit into two blocks, each lasting about 7-8 minutes (24 presentations spaced by roughly 20s each), so that the first block covers the first phase while the second block covers the second phase. In Figure 4C we show the spectrograms of scopolamine (50 mM) infusion followed by odor presentation sessions in 4 rats to confirm that the first odor block (green line) and second odor block (red line) bracketed this transition. We used the same odor block design for all sessions to avoid introducing new variables, but since APV did not show biphasic effects we combined the APV data from both blocks.

While the scopolamine and APV effects were fairly consistent from session to session, the oxotremorine effects were highly variable. In the early sessions without odor presentations, oxotremorine tended to increase gamma power (Fig. 3Cii, gray trace), but in 70% of the odor presentation sessions oxotremorine reduced gamma power (Fig. 3Cii, pink trace). In those sessions, gamma suppression was followed by a rebound gamma increase, similar to the scopolamine sessions. Because of this, we used the same odor block design for oxotremorine as scopolamine. Oxotremorine is known to produce excitatory and inhibitory effects in the OB through differential activation of M1 and M2 receptors (Smith et al., 2015, see *Discussion*), and this may in part be responsible for the highly variable effects on gamma. We include oxotremorine data here for completeness, but we admit that more work must be done to dissect its effect on OB oscillations.

### Muscarinic receptors

Our previous modeling work (Osinski and Kay, 2016) predicted that reducing GC excitability would lift the frequency of LFP oscillations induced by strong odors out of the beta regime into the low gamma range but would leave LFP oscillations evoked by weak odors in the beta regime with possibly some increase in power (Fig. 1B). To test these effects, we infused scopolamine (38 mM & 50 mM), a nonselective muscarinic antagonist with twice the affinity for M2 as for M1 receptors (Bolden et al., 1992), through a cannula positioned in the GC layer (Fig. 2D). Muscarinic drugs are known to modulate GC excitability in a complex manner (Castillo et al., 1999; Mandairon et al., 2006; Pressler et al., 2007; Devore and Linster, 2012; Li and Cleland, 2013; Smith et al., 2015). With scopolamine, our intention was to prevent heightened excitability states from occurring, thus placing GCs in an impaired excitability state, as described in the modeling results in Figure 1B. We chose one high volatility odorant, ethyl-2-methylbutyrate (EMB), and one low volatility odorant, geraniol (GER), to probe the effects induced by strong and weak odors, respectively. As described earlier (Fig. 4), we divided odor presentation sessions into two blocks of 24 trials each lasting ~7-8 minutes to cover the biphasic scopolamine effect on the background LFP.

#### Results from a single scopolamine session

We first take a close look at a representative scopolamine (50 mM) session and its associated saline session from a single rat (Fig. 5), and then we examine the summary statistics of all the scopolamine sessions across all rats (Fig. 6). Figure 5A shows single trials of EMB- and GER-evoked oscillations in LaOB, LmOB (infusion site), RmOB, and LaPC after saline and scopolamine (50 mM) infusions. In these sessions, EMB was presented in the 1^st^ odor block and GER in the 2^nd^. EMB evoked prominent beta oscillations on all channels after saline infusion (labeled β, Fig 5Ai). LaOB beta was typically smaller in amplitude than LmOB, consistent with other recordings in our laboratory, possibly because anterior OB receives fewer feedback fibers than more posterior parts or simply because the increased curvature in the anterior end of the bulb reduces the coherence of the laminar cortical field. After scopolamine 50 mM infusion, the EMB-evoked LaOB and LmOB beta oscillations were abolished and appeared to be replaced with a high power low gamma frequency oscillation (labeled low γ, Fig. 5Aiii). The contralateral (RmOB) beta oscillation, where no drug was delivered, was still present (labeled β, Fig 5Aiii). GER did not evoke visible beta oscillations under the saline condition (Fig 5Aii), but did evoke visible beta oscillations after scopolamine infusion (Fig 5Aiv). In these trials the apparent opposite effect of scopolamine on EMB- and GER-evoked beta oscillations followed our model predictions of a bidirectional effect (Fig. 1B).

**Figure 5.**
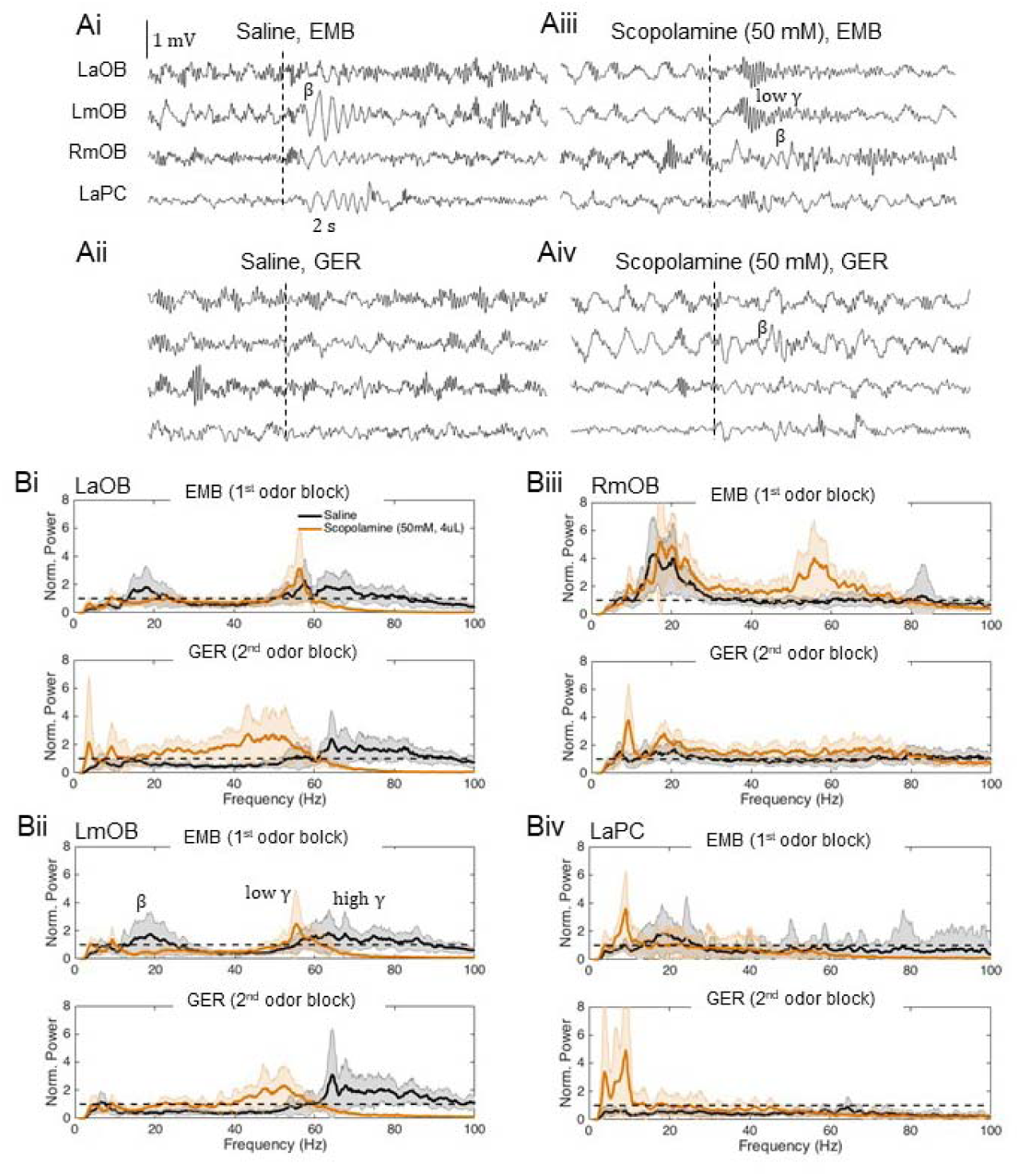
Representative scopolamine session and associated saline session from a single rat. **A:** Representative LFP traces during EMB (top) and GER (bottom) presentations in LaOB, LmOB, RmOB, and LaPC after saline (left) and scopolamine (right) infusions from a single rat. Twenty such trials are used to produce the power spectra in B. Odor presentation times are indicated by dashed vertical lines. In this session the EMB-induced beta in the LOB (labeled β) was visibly abolished and appeared to be replaced by a low gamma oscillation (labeled low γ). This occurred in 9 out of 12 of the scopolamine (50 mM) sessions and 4 out of 12 of the scopolamine (38 mM) sessions across the 6 rats. Beta persisted in the RmOB after scopolamine infusion. **B:** Mean power spectra of LaOB (Bi), LmOB (Bii), RmOB (Biii), and LaPC (Biv). For each brain region we show the LFP power spectra during EMB (top) and GER (bottom) presentation after saline (black) and scopolamine (orange) infusion. Power was normalized by the pre-infusion baseline, represented by the horizontal dashed line at 1. In the LmOB EMB spectrum (Bii, top) we annotate the beta (β), low gamma (low γ), and high gamma peaks (high γ).

**Figure 6.**
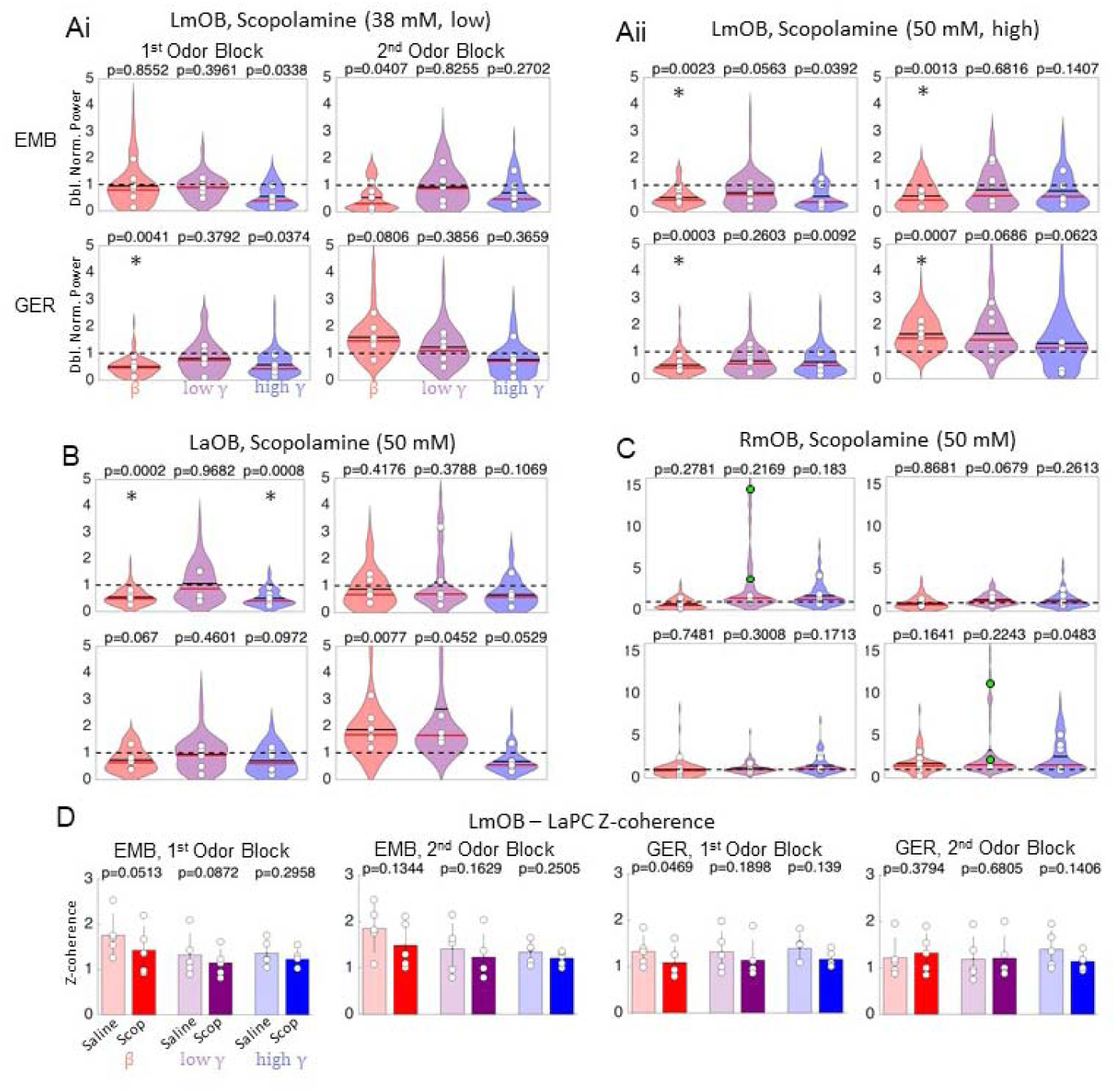
Scopolamine effects on odor induced LFP oscillations. **A:** Violin plots summarizing effects of low (38 mM, Ai) and high (50 mM, Aii) scopolamine doses on LmOB (infusion site) in all 6 rats for presentation of EMB (top) and GER (bottom) during 1^st^ and 2^nd^ odor blocks. White circles represent means of each rat. Power for each session was doubly normalized, first by baseline and then by its closest saline session (black dashed line). A value of 1 indicates no difference from the saline condition. Black solid horizontal lines on the violins show means for all rats; red horizontal lines are medians. Asterisks represent p < 0.004 for the post hoc comparisons between mean saline and drug for each rat. **B, C:** Violin plots summarizing effects of scopolamine (50 mM) on LaOB and RmOB (same annotations as A). Although the scopolamine effect in RmOB was not significant in the *post hoc* comparison across all rats, two rats showed very strong low gamma power (colored green for visualization). Note that the y-scale is increased for RmOB to fit the huge increase in low gamma power that occurred in one rat. D: Z-coherence between LmOB and LaPC under saline and scopolamine (50 mM) conditions was computed for EMB and GER in both odor blocks.

Figure 5B shows the average power spectra (normalized by baseline) of the single sessions from which the representative trials in Figure 5A were taken. The average power in the beta frequency range for LaOB and LmOB during the 1^st^ odor block (EMB) is noticeably reduced, and a peak in the low gamma frequency band appears after scopolamine infusion (Fig 5Bi,ii, top; peak at ~55 Hz). This low gamma peak represents the low gamma oscillation that appears to occur in place of the beta oscillation, seen in the LFP trace in Fig 4Bii. There is also an increase in broadband low gamma in the 2^nd^ odor block (GER). Unlike the 1^st^ odor block however, this increase in low gamma is most likely not odor-evoked, but a consequence of the rebound gamma increase that occurs in the later phase of the scopolamine effect as shown in Figures 3 & 4. The distinction between the gamma frequencies in the EMB- and GER-induced LFP spectra may be related to the two different gamma subtypes, gamma1 and gamma2, noted by Kay (2003) (see *Discussion*). There was a small increase in GER-evoked beta under scopolamine in the representative trial shown in Figure 5Aiv, and, on average, the GER-evoked beta power was slightly elevated relative to baseline for this session (Fig. 5Bi,ii, bottom). The LaPC showed negligible effects of EMB-evoked (Fig. 5Biv, top) and GER-evoked (Fig. 5Biv, bottom) beta following scopolamine infusion. Because PC tends to produce beta frequency activity spontaneously (Poo and Isaacson, 2009), the baseline normalized LaPC beta was fairly low even when there was a visibly evoked beta oscillation. All three regions in the left hemisphere show a persistent decrease in high gamma oscillations lasting through both phases of the scopolamine effect. Although this occurred to some degree in 4 of 6 rats, the effect overall was not significant (see Fig. 6). In the RmOB there was no effect on beta power but a surprising increase in low gamma power following scopolamine infusion in this rat (Fig. 5Biii). There were prominent contralateral effects in approximately half of the rats, and this is discussed in more detail later (see Fig. 6C). We present this representative data set from a single rat because the replacement of beta by low gamma oscillations following scopolamine infusion could only be inferred by looking at the individual LFP traces themselves (comparing Fig. 5Bi and Fig. 5Biii), and not from summary statistics.

#### Scopolamine effects in the OB

We now turn to address drug effects across all rats for low and high doses of scopolamine on the beta (15 – 30 Hz), low gamma (40 – 60 Hz), and high gamma (60 – 100 Hz) frequency bands. In order to put power values from different rats on the same scale we normalized the LFP power by both pre-infusion baseline and saline power (see *Methods*). 5-way ANOVAs were computed, with subject, drug treatment (saline vs. drug), odor, frequency band, and block (early vs. late) as factors for each electrode location and drug concentration. The results of post hoc t-tests between the means of the saline and drug sessions for each rat and electrode location are reported in Fig. 6. As described in *Methods*, the Bonferroni correction on post hoc tests sets the significance threshold at p < 0.004 for drugs that showed a biphasic effect. For these and all subsequent descriptions of the drug effects, we first present the results of our ANOVA analyses, including the effects of all factors. After that, for electrode locations that show significant main effects or interactions involving the drug treatment, we present post hoc comparisons. Other effects (odor, subject, frequency band) have been extensively covered in our previous reports (Lowry and Kay, 2007; Kay et al., 2009; Kay, 2014; Frederick et al., 2016).

Effects on the left OB (treatment side) were similar across the two drug concentrations. For the low concentration (38 mM) scopolamine treatment, we found significant main effects of all five factors on LFP power in both sites in the OB (LmOB: subject p=7.4e-8, η^2^ 0.16; drug p=0.0019, η^2^ 0.03; odor p=1.03e-7, η^2^ 0.11; block p=0.0076, η^2^ 0.0003; frequency band p=7.77e-5, η^2^ 0.07; LaOB: subject p=0.0324, η^2^ 0.05; drug p=0.0389, η^2^ 0.02; odor p=1.00e-6, η^2^ 0.12; block p=0.0434, η^2^ 0.003; frequency band p=0.00001, η^2^ 0.08). Some of these effects were expected because of our prior knowledge about how the volatility of odors affects the power of beta oscillations (odor, band) and differences in electrode placement and therefore power of different oscillation frequencies across subjects (subject). Importantly, there was also a significant interaction between drug and odor at the injection site only (LmOB: p=0.0366, η^2^ 0.15), plus additional interactions at that site between subject and odor (p=0.0249, η^2^ 0.04), subject and frequency band (p=8.76e-7, η^2^ 0.17), odor and block (p=0.0447, η^2^ 0.01), and odor and frequency band (p=2.66e-6, η^2^ 0.10). Significant interaction effects for the LaOB were subject x band (p=2.04e-6, η^2^ 0.22), odor x block (p=0.005, η^2^ 0.03), and odor x band (p=0.0007, η^2^ 0.07).

For the high concentration (50 mM) scopolamine treatment, we again found significant main effects for all five factors at the injection site (LmOB: subject p=1.93e-7, η^2^ 0.15; drug p=0.0020, η^2^ 0.03; odor p=2.57e-7, η^2^ 0.09; block p=0.0114, η^2^ 0.02; frequency band p=5.24e-5, η^2^ 0.07). We also found significant interactions between drug treatment and odor (p=0.0043, η^2^ 0.03) and drug treatment and block (p=0.0211, η^2^ 0.02), in addition to expected subject x band (p=1.93e-5, η^2^ 0.14) and odor x band (p=1.45e-6, η^2^ 0.09) effects. At the anterior OB recording site we did not find significant main effects of subject or odor, but we did find significant main effects of drug, block and frequency band (LaOB: drug p=0.0087, η^2^ 0.0001; block p=0.0327, η^2^ 0.004; band p=2.58e-5, η^2^ 0.11). At this site, we also found significant interaction terms involving drug treatment (drug x odor, p= 0.0154, η^2^ 0.03; drug x block, p=0.0066, η^2^ 0.03; drug x band, p=0.0031, η^2^ 0.03), plus additional interactions (subject x band, p=0.0006, η^2^ 0.16, and odor x band, p=0.0043, η^2^ 0.05) as expected.

*Post hoc* comparisons of drug vs. saline effects for each electrode site and frequency band shed more light on main effects and interactions. In the first odor block, LmOB beta power was significantly reduced for the high volatility odorant (EMB) only for the high scopolamine dose (50 μM, Fig. 6Aii top) but was significantly reduced for both low and high doses when the low volatility odorant (GER) was presented (Fig. 6Ai,ii bottom). It is possible that the strong beta-evoking tendency of EMB may be counteracting the beta suppressing effects of scopolamine only at the low dose. GER evokes beta oscillations more weakly, and therefore the effect of scopolamine dominates even in the low dose.

In the second odor block, effects on beta power diverged between the strong and weak odors, confirming the bidirectional effect predicted by our model. For the high scopolamine dose in the second block (Fig. 6Aii), EMB-evoked beta was suppressed just as it was in the 1^st^ odor block, but GER-evoked beta power was significantly increased. Similar trends are seen for the low scopolamine dose, though they did not pass our threshold for significance. The LaOB LFP power followed a pattern similar to LmOB, but only the reduction of beta in the 1^st^ odor block was significant (Fig. 6B). The weaker effects in LaOB may be attributed to weaker overall beta signals in the anterior OB or possibly insufficient spread of the drug in some rats (Fig. 2Bi). The complete replacement of EMB-evoked beta by low gamma following scopolamine (50 mM) infusion (as seen in the single trials of Fig. 5A) was seen in 4 out of 12 scopolamine (38 mM) sessions and 9 out of 12 scopolamine (50 mM) sessions (from both odor blocks), while in the sessions beta amplitude was either reduced or unaffected. The replacement of beta by low gamma can be inferred only from viewing the odor-evoked periods, not from the power over the entire 4s trial periods.

There were no significant effects of either low or high doses of scopolamine on the LmOB gamma bands during the odor exposure periods, though there was a tendency for high gamma to be suppressed in during the 1^st^ odor block, consistent with gamma suppression during the early phase of scopolamine in the no odor condition (Fig. 3). There was a significant reduction in LaOB high gamma power for EMB in the 1^st^ odor block (Fig. 6B). Four of the 6 rats showed enhanced low gamma power in the 2^nd^ odor block of high dose scopolamine, presumably due to the rebound low gamma intensification described in Figures 3 & 4. One possible reason why rebound gamma did not show a significant gamma increase in all rats is that odorants tend to increase beta while suppressing gamma (Buonviso et al., 2003).

#### Scopolamine effects in contralateral OB

As touched upon in Figure 5Bii, we noticed some strong but inconsistent contralateral effects in the gamma frequency bands following scopolamine infusion into the LmOB. The sessions of two rats in particular (both sessions where EMB was presented in block 1) had extremely high gamma power under scopolamine, especially in the low gamma band. The means of these two rats are colored green in the low gamma power bands of Figure 6C for the EMB 1st and GER 2nd odor blocks (same session). These are the same two rats that had the furthest posterior spread of dye in post-mortem inspection (two green lines in Fig 2C).

The results of the ANOVAs at the two different scopolamine concentrations showed no main effect of drug treatment on this side, but did show other significant main effects, as would be expected (38 mM scopolamine: odor, p=1.65e-6, η^2^ 0.12; band, p=7.63e-9, η^2^ 0.21; 50 mM scopolamine: odor, p=0.007, η^2^ 0.04; band, p=0.0118, η^2^ 0.05). At both concentrations there was a significant interaction between drug and frequency band (38 mM: p=0.0025, η^2^ 0.06; 50 mM: p=0.0015, η^2^ 0.08), which is likely driven by the increase in low gamma power. We also found a significant odor x band interaction (38 mM: p=0.4.25e-7, η^2^ 0.18; 50 mM: p=0.0170, η^2^ 0.05).

The increase in low gamma power was not odor evoked, but persisted between odor presentations and lasted throughout the EMB and GER odor blocks. While the drug x band interaction was significant with a medium effect size, *post hoc* t-tests were not significant because of the wide variance across subjects. When the increase in low frequency gamma did occur, it was obvious to the eye, even during data acquisition. We suspect that in the sessions where there was a contralateral effect, some of the drug spread posteriorly into the anterior olfactory nucleus (AON). Contralateral projections through the anterior commissure linking the left AON to the right OB are known to exist (Nickell and Shipley, 1993), and it is possible that blocking muscarinic modulation of these fibers is driving the contralateral increase in gamma.

#### Scopolamine effects in LaPC

In addition to LOB and ROB activity, LaPC activity was also recorded in each rat (Fig. 5A, bottom trace). One rat’s LaPC data had to be discarded due to poor quality signals.

The results of the ANOVAs showed the same effects for both drug doses. There were no significant effects of drug treatment, either as main effects or interactions (LaPC 38 mM: subject, p=2.23e-10, η^2^ 0.22; odor, p=4.25e-7, η^2^ 0.09; band, p=9.68e-9, η^2^ 0.14; remaining NS. LaPC 50 mM: subject, p=3.29e-10, η^2^ 0.20; odor, p=1.55e-5, η^2^ 0.06; band, p=5.40e-10, η^2^ 0.16; remaining NS.). There were expected interactions that depend on differences across subjects and odors: subject x odor (38 mM: p=9.96e-6, η^2^ 0.09; 50 mM: p=0.0014, η^2^ 0.06), subject x band (38 mM: p=3.67e-8, η^2^ 0.18; 50 mM: p=1.31e-6, η^2^ 0.15), and odor x band (38 mM: p=5.53e-7, η^2^ 0.10; p=1.82e-6, η^2^ 0.09). These results show that there was no effect on aPC power in any of the frequency bands due to scopolamine action in the OB.

We had expected that a reduction in OB beta power by scopolamine would also reduce ipsilateral aPC beta power, because beta oscillations require intact bidirectional OB-PC connections (Neville and Haberly, 2003; Martin et al., 2006), and GCs receive most of their cortical inputs from the ipsilateral aPC. The results suggest that beta oscillations in the PC can be generated with some degree of independence from the OB, even though they still require intact projections from the bulb (at least under anesthesia, Neville and Haberly, 2003). This stability could be attributed to the PC’s own tendency to generate beta frequency oscillations in response to odor stimulation. Indeed, a study by Poo and Isaacson (2009) found prominent PC beta oscillations in urethane-anesthetized rats, even when anesthesia depresses the feedback inputs into the OB.

We also computed the Z-coherence between OB and PC for scopolamine (50 mM) sessions and the associated saline sessions. A 5-way ANOVA showed significant main effects of subject, drug, odor, and frequency band (subject, p=1.44e-33, η^2^ 0.62; drug, p=0.0003, η^2^ 0.05; odor, p=3.61e-7, η^2^ 0.04; band, p=1.14e-6, η^2^ 0.04; block NS). The drug effect was different across subjects (subject x drug, p=0.0066, η^2^ 0.02), and there were other significant interactions not involving scopolamine treatment (subject x band, p=8.22e-6, η^2^ 0.06; odor x band, p=4.8e-7, η^2^ 0.04; remainder NS). While there was a significant effect of scopolamine overall as a slight reduction of coherence, none of the post hoc comparisons were significant (Fig. 6D). Thus, OB-PC beta coherence was still high even when beta was seemingly eliminated in the OB. This suggests that a small component at beta frequency persisted in the OB, even when beta power was suppressed to baseline levels by scopolamine, and that this is enough to support beta band coherence with the aPC.

#### Oxotremorine effects

To complement the antagonistic effects of scopolamine, we also infused a muscarinic agonist into the OB. We chose oxotremorine, because it has already been used in several studies of bulbar cholinergic modulation (Mandairon et al., 2006; Smith et al., 2015). We expected oxotremorine and scopolamine to have opposite effects, but oxotremorine produced inconsistent results (Fig. 3). While the results are somewhat equivocal, we include them here for completeness.

Results from our ANOVA analysis show that all factors show significant main effects at the injection site (LmOB: subject, p=0.0002, η^2^ 0.11; drug, p=0.0033, η^2^ 0.02; odor, p=1.46e-5, η^2^ 0.09; block, p=0.0173, η^2^ 0.02; band, p=0.0054, η^2^ 0.05). There were no significant interactions of any of the other factors with drug, but there were some other significant interactions: subject x band (p=0.0131, η^2^ 0.09), odor x block (p=0.0013, η^2^ 0.05), and odor x band (p=3.86e-5, η^2^ 0.05). At the anterior OB site, there were main effects only of drug and odor (LaOB: drug, p=0.0019, η^2^ 0.06; odor, p=0.0028, η^2^ 0.06), no significant interactions with drug effects, but other significant interactions (odor x band, p=0.048, η^2^ 0.04; subject x band: p=0.0122, η^2^ 0.12; odor x band, p=0.0021, η^2^ 0.08).

As opposed to scopolamine, oxotremorine did significantly affect PC activity. We found significant main effects for all factors except odor block (LaPC: subject, p=0.0003, η^2^ 0.11; drug, p=0.0156, η^2^ 0.03; odor, p=0.0007, η^2^ 0.06; band, p=0.0002, η^2^ 0.10). There were no significant interactions with drug effects, but there were interactions between subject and odor (p=0.0360, η^2^ 0.04), subject and frequency band (p=0.0009, η^2^ 0.13), odor and block (p=0.0282, η^2^ 0.02), and odor and frequency band (p=0.0003, η^2^ 0.09). The contralateral OB also showed significant main effects, including a drug effect (RmOB: drug, p=0.0226, η^2^ 0.02; odor, p=1.70e-5, η^2^ 0.09; band, p=6.28e-11, η^2^ 0.25, remaining factors NS). No factors showed significant interactions with drug, but other interactions were significant: odor x block (p=0.0139, η^2^ 0.03) and odor x band (p=1.91e-7, η^2^ 0.15).

These effects played out in the *post hoc* analysis in somewhat confusing ways. Following oxotremorine (30 μM) infusion into LmOB, there was a significant decrease in LaOB beta power during presentation of GER in the 2^nd^ block (Fig. 7A) and LmOB beta power during EMB in the 1^st^ odor block (Fig. 7B). This was surprising, because scopolamine also blocked beta in the 1^st^ EMB odor block, but increased beta in the 2^nd^ GER odor block (Fig. 6Aii). So, the two drugs produced the same effect in the first block but opposite effects on GER in the second block.

**Figure 7.**
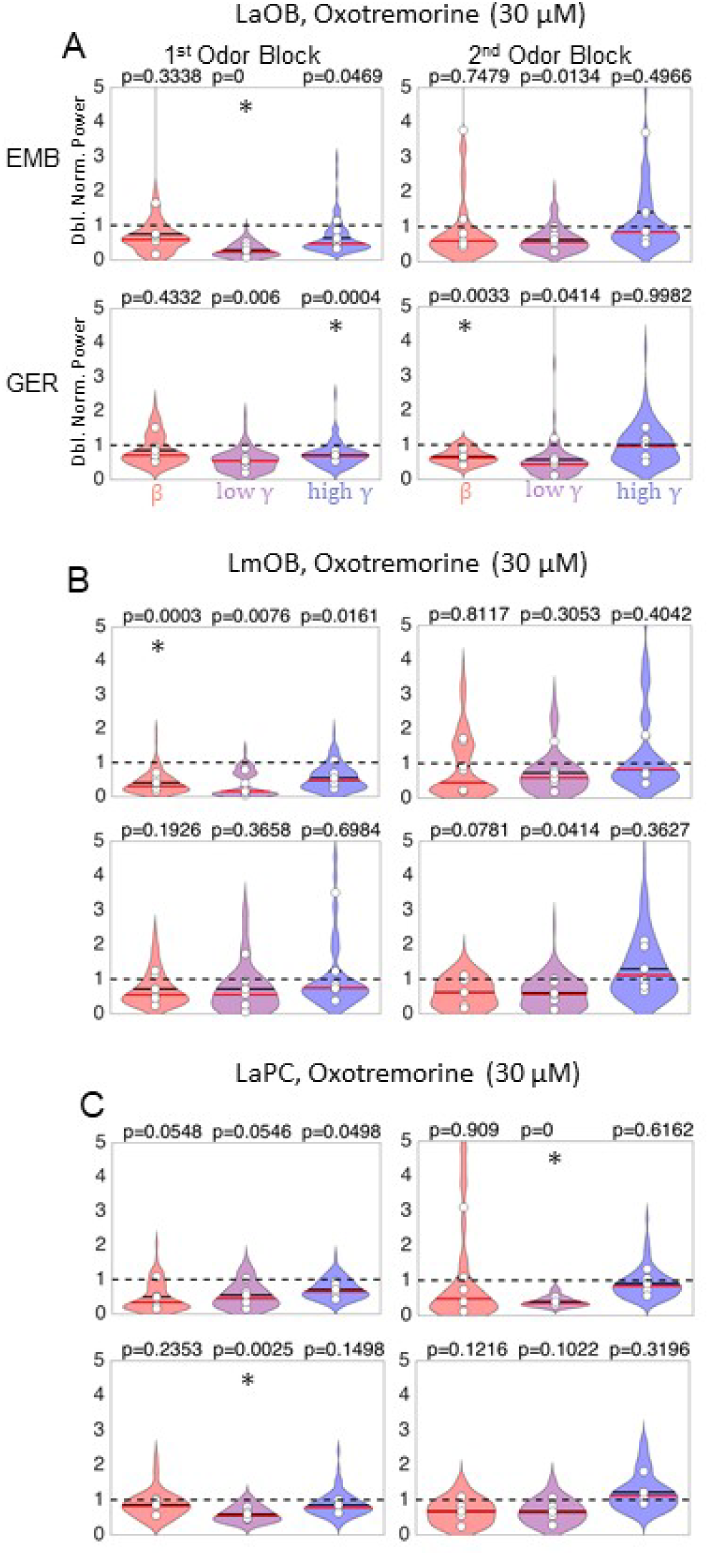
Oxotremorine effects on odor-induced LFP oscillations. Violin plots summarizing the effects of oxotremorine on LaOB (A), LmOB (B, infusion site), and LaPC (C) during presentation of EMB (top) and GER (bottom). (There were no significant *post hoc* drug effect comparisons for RmOB.) White circles represent means of each rat. (Normalization, color codes, and significance asterisks are the same as in Fig. 6).

However, while scopolamine may have blocked EMB-induced beta by reducing GC GABA release to the point that oscillations could not be sustained, oxotremorine may have blocked EMB-induced beta by driving excessive GABA release, tipping the balance in the other direction where inhibition dampens excitation too much. Such an effect was also seen in our model, where oscillations were only sustained in a range where there was a sufficient balance of excitation and inhibition. It is also possible that oxotremorine may inhibit GCs through action on M2 receptors or inhibit MCs themselves (Smith et al., 2015).

LaOB low gamma was significantly reduced in the 1^st^ EMB odor block (close to significance in LmOB), and high gamma was reduced in the 1^st^ GER odor block. As noted in Figure 3, oxotremorine tended to increase gamma power when no odor was present, but suppress it when odor was present. Interestingly, gamma suppression in the LaPC was significant both for the 2^nd^ odor block EMB presentations and 1^st^ odor block GER presentations (Fig. 7D). *Post hoc* analyses of RmOB responses to oxotremorine infusion into LmOB showed no significant comparisons between saline and drug. Because of the inconsistent effects of oxotremorine (Fig. 3 Cii) and its likely non-specific effects (see *Discussion*) we refrain from making any strong conclusions about the effects of oxotremorine on beta oscillation generation in the OB.

### NMDA receptors

#### APV effects

We also tested another prediction of our model, that beta oscillations can be sustained independently of NMDAR currents, but that NMDAR currents are critical to sustain gamma oscillations. (In our model, beta oscillations rely critically on N-Type mediated Ca^2+^ currents.) We tested this prediction by infusing APV (100 μM), a selective NMDAR antagonist, into the LmOB. Because APV had a uniform effect lasting the entire session (Fig. 3Ciii), we combined data from the 1^st^ and 2^nd^ odor blocks, the results of which are summarized in Figure 8.

**Figure 8.**
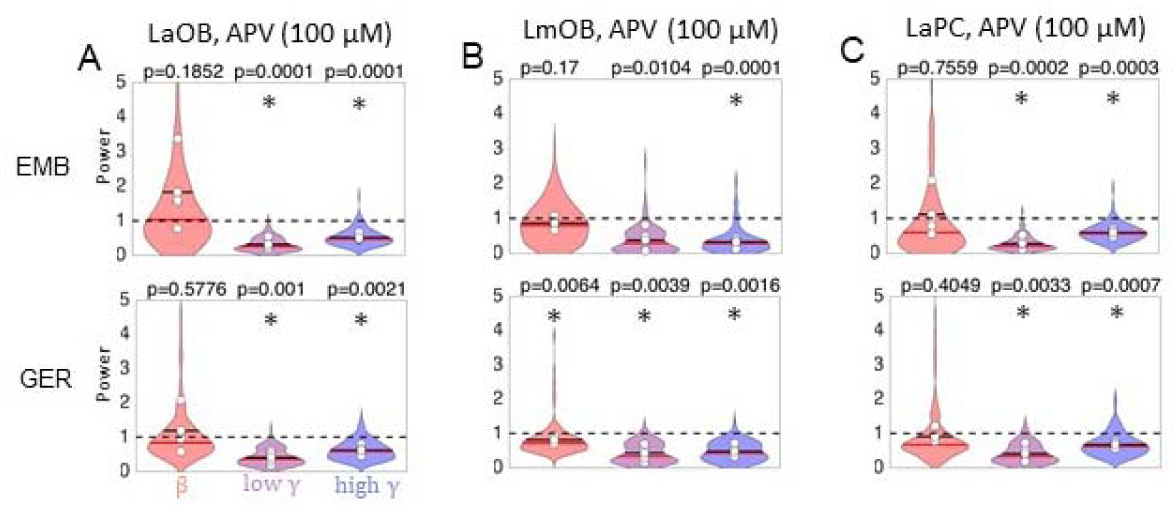
APV effects on odor-induced LFP oscillations. **A:** Violin plots summarizing effects of APV infusion into LmOB on LaOB in all 6 rats for presentation of EMB (top) and GER (bottom). 1^st^ and 2^nd^ odor blocks were combined for analysis of APV data (Fig. 3). (Symbols and normalization are the same as in Figs. 6 and 7.) Asterisks represent p < 0.008 (adjusted for 6 comparisons) for the post hoc T-tests between mean saline and drug for each rat. **B, C:** Violin plots summarizing effects of APV in LmOB (infusion site) and LaPC (same annotations as A). APV strongly suppresses gamma in all left hemisphere locations.

We analyzed the data with a 4-way ANOVA to test the influence of subject, drug, odor, and frequency band. At the injection site, all four factors showed significant main effects (LmOB: subject, p=0.0032, η^2^ 0.11; drug, p=0.0009, η^2^ 0.09; odor: p=2.98e-5, η^2^ 0.16; band, p=0.0002, η^2^ 0.15). APV treatment did not show significant interactions with any other factors at this site (subject x band, p=0.0137, η^2^ 0.13; odor x band, p=0.0004, η^2^ 0.13; remaining interactions NS). The main effects were somewhat different at the anterior OB site (LaOB: subject, NS; drug, p=0.0077, η^2^ 0.02; odor, p=8.93e-5, η^2^ 0.11; band, p=1.19e-7, η^2^ 0.34). APV treatment did show a significant interaction with frequency band at this site (p=0.0021, η^2^ 0.08), and there were some other interactions (subject x band, p=0.0337, η^2^ 0.08; odor x band, p=2.64e-5, η^2^ 0.17; other interactions NS).

As predicted by our model and in agreement with studies in mice (Lepousez and Lledo, 2013), post hoc tests showed that LaOB and LmOB gamma oscillations were almost completely abolished (Fig. 8A,B). LaOB and LmOB beta power was largely unaffected, except for the LmOB GER presentations, which showed a small but significant decrease. It is possible that the decrease in LmOB beta for GER presentations reflects a stimulus dependent effect, similar to the contrasting effects of EMB and GER seen during scopolamine infusion. However, we did not address this scenario in the model.

Analysis of the contralateral OB data showed no main or interaction effect of APV treatment (RmOB: odor, p=3.41e-5, η^2^ 0.11; band, p=1.51e-8, η^2^ 0.38; subject x band, p=0.0093, η^2^ 0.10; odor x band, 1.79e-6, η^2^ 0.22; remainder NS). We therefore omit the post hoc analysis. The ipsilateral aPC showed significant main effects for all factors (LaPC: subject, p=0.0011, η^2^ 0.13; drug, p=0.0344, η^2^ 0.03; odor, 0.0003, η^2^ 0.10; band, 2.01e-5, η^2^ 0.20). Interaction effects did not involve APV (subject x odor, p=0.0369, η^2^ 0.06; subject x band, p=0.0031, η^2^ 0.16; odor x band, p=0.0002, η^2^ 0.14; remainder NS).

*Post hoc* tests show that in contrast to the relative insensitivity of LaPC beta oscillations to scopolamine infusion into the LmOB, we found strong gamma suppression in LaPC after APV infusions into the LmOB (Fig. 8D), mirroring the effects in the OB.

## Discussion

We tested two predictions of our previously published computational model (Osinski and Kay, 2016b; Fig 1): 1) beta oscillations are produced under conditions of heightened GC excitability, 2) beta oscillations can be generated independently of NMDA currents, while gamma oscillations cannot. We tested the first prediction by infusing muscarinic drugs into the OB (Figs. 3-7), and the second prediction by infusing APV, a selective NMDA antagonist (Figs. 3 & 8).

We found that scopolamine (50 mM) reduced EMB- and GER-evoked beta oscillations in the 1^st^ odor block, but after a few minutes (in the 2^nd^ odor block) we observed a divergent effect, where EMB-induced beta was suppressed and GER-induced beta was enhanced (Fig. 5A). The divergent effect in the 2^nd^ odor block also aligns well with our model’s predictions, that reducing GC excitation in strong stimulus input regimes (EMB) would decrease beta power and in weak stimulus input regimes (GER) would increase beta power. In 9 out of 12 of the scopolamine (50 mM) sessions, the EMB-evoked beta oscillation was replaced by a low gamma oscillation (Figs. 4A, 5Aii), as our model predicted for strong odors (green square Fig. 1B). The results of APV (100 μM) infusions also closely followed our model predictions (Fig. 7), knocking out gamma, but not beta, oscillations. However, oxotremorine (30 μM) effects were more difficult to interpret (Fig. 2 & 6). By also recording in LaPC and RmOB we were able to investigate the wider network that supports and is impacted by these oscillations across brain regions beyond the scope of the model. We found that PC beta oscillations were relatively insensitive to changes in OB beta power, but PC gamma oscillations were much more sensitive (Fig. 5D & 7D).

### Complexity of muscarinic drug effects

Many studies of cholinergic modulation in the OB treat muscarinic receptors as modulators of GC excitability (Nickell and Shipley, 1993; Castillo et al., 1999; Martin et al., 2006; Pressler et al., 2007; Devore and Linster, 2012; de Almeida et al., 2013; Li and Cleland, 2013). Together these studies give the impression that bulbar muscarinic receptors are exclusively involved in regulation of GC excitability, and this is indeed why we chose to use muscarinic drugs in this study. Although M1 and M2 receptors are found in highest density in the GC layer and external plexiform layers (where GCs form dendrodendritic synapses with MC lateral dendrites) they can be found throughout the OB (see Allen Mouse Brain Atlas CHRM1 & CHRM2 genes online; Fonseca et al., 1991; Lein et al., 2007). Therefore, our infusions of scopolamine are mostly targeting GCs and GC dendrites but to a lesser degree also targeting other cells as well.

Oxotremorine has been shown to have strong inhibitory effects on GCs and MCs in the main OB and excitatory effects on MCs and GCs in the accessory OB (AOB) through differential activation of M1 and M2 receptors (Smith and Araneda, 2010; Smith et al., 2015). Interestingly, Smith and Araneda also hypothesized that the direction of the effect may depend on the strength of input to GCs. They argued that when excitatory input onto GCs is weak, M2-mediated hyperpolarization is the predominant effect, reducing inhibition of MCs, but when excitatory input onto GCs is strong (*i.e*., from excited MCs), the M1-mediated afterdepolarization will prevail, prolonging GC activation and increasing MC inhibition. There may be an interaction between oxotremorine effects on muscarinic receptors and strong odor stimulation triggering long lasting depolarizations in GCs (Egger et al., 2005; Pressler et al., 2007). These competing effects may explain the observed tendency for oxotremorine to increase gamma power when odors were absent, but decrease gamma when odors were present (Figs. 3Cii, 7).

Besides having opposite effects on M1 and M2 receptors, oxotremorine is also capable of activating nicotinic receptors, even though it is classified as a muscarinic agonist (Akk et al., 2005). Given the complexity of interactions, it is not altogether surprising that a drug as nonspecific as oxotremorine produces messy effects in an awake behaving animal, which itself is also most certainly releasing endogenous ACh during the recording session.

Scopolamine produced a biphasic effect in the LFP, first reducing broadband gamma power for ~7 min, followed by an intensification of low gamma while high gamma was still suppressed (Figs 3 & 4). The opposite effects on the two gamma bands in the 2^nd^ phase may reflect the distinction between type 1 gamma, which depends on MC-GC interactions, and type 2 gamma, which may depend on GABAergic inhibition of GCs, as reported by Kay (2003). Although we ignored inhibition of GCs in our model for simplicity, it is quite possible that in reality scopolamine influences inhibition of GCs to produce contrasting effects on gamma 1 and gamma 2.

### Spatial extent of gamma and beta oscillations

Gamma oscillations are thought to be more spatially localized to individual cortical areas and even parts of these areas, while beta oscillations are thought to represent a coordinated oscillation supported by bidirectional connections between OB and PC (Martin et al., 2006; Kay et al., 2009; Kay and Lazzara, 2010). Curiously, we found that aPC *beta* power was not significantly reduced when OB beta was reduced by scopolamine, but aPC *gamma* was significantly reduced when ipsilateral OB gamma was suppressed by infusion of APV (Fig. 8D). It has long been known that oscillatory evoked potentials in PC depend on but are not driven by OB input (Freeman, 1968), and APV desynchronizes MCs from population gamma coordination but does not suppress their firing. Thus, our results suggest that this input must be coordinated with the gamma rhythm in order for the PC to be capable of generating gamma spontaneously. On the other hand, this suggests that PC beta oscillations do not require beta band synchronized inputs from MCs, just some level of tonic excitation, or that the PC does not require very much beta band input from the OB (OB-aPC coherence was hardly affected under scopolamine). Future studies should address the mechanisms for beta vs. gamma generation in the aPC in order to understand this interesting dichotomy.

The effect of APV on anterior OB is very similar to that on medial OB, but with even more aggressive suppression of both gamma frequency bands (Fig. 8A,B). Although in some of the rats the drug did not appear to spread fully to the LaOB (see Fig. 2B), the gamma suppression was seen in all rats, suggesting that GCs in the more posterior bulb influence the activity of MCs in the anterior portion of the bulb. This aligns well with studies that have shown GCs to mediate between distant MCs via the long MC lateral dendrites, which can span the bulb (Migliore and Shepherd, 2008).

### Contralateral effects

In some of the rats there were strong contralateral effects following scopolamine (Fig. 6C) and APV (Fig. 8C) infusions. We assume that these effects were not caused by drug spreading into the RmOB, since dye spread was localized to the left side in all rats (Fig. 5E). Instead, we suspect that these effects were caused by drug spreading into the anterior olfactory nucleus (AON), which is innervated by fibers that link the two hemispheres through the anterior commissure. Indeed, those rats that had the most posterior dye spread (Fig. 2C) had the largest contralateral effects. Muscarinic modulation of anterior commissure fibers was reported by Nickell and Shipley (1993), who hypothesized that cholinergic inputs to MOB may modulate cross-bulbar information. The AON is known to densely innervate GCs (Price and Powell, 1970). Therefore, cross-bulbar communication may be regulated by fibers controlling GC excitability in both hemispheres. We are not aware of any studies relating NMDA-dependent modulation of cross-bulbar information, but our study suggests that this may also be possible. Though it was not our intention, our study provides motivation to more thoroughly study cross-bulbar mediation of gamma oscillations through cholinergic and NMDA dependent effects in the AON.

### Implications for odor discrimination

Bulbar infusions of scopolamine have been shown to reduce spontaneous odor discrimination of closely related odorants (fine odor discrimination) in the absence of reinforcement learning (Mandairon et al., 2006; Chaudhury et al., 2009). Gamma and gamma-like oscillations have been functionally related to fine odor discrimination in honeybees, mice, and rats (Stopfer et al., 1997; Nusser et al., 2001; Beshel et al., 2007). Therefore, it might be expected that bulbar scopolamine would reduce gamma oscillation power. Our model predicted, and we found, that under scopolamine instead of gamma, beta oscillations were reduced or increased during strong or weak odor exposure, respectively. (There was some suppression of high gamma in the LaOB, but only in response to EMB and only in the first block.)Because the same conditions that increase odor generalization (muscarinic block) also manipulate beta oscillations, it is possible that beta oscillations are also involved in fine odor discrimination. Thus far, however, the evidence does not support this inference, at least in the context of reinforcement learning, where beta oscillations are either not increased or suppressed during fine odor discrimination (Beshel et al., 2007; Frederick et al., 2016). However, it should be noted that muscarinic antagonist effects on odor perception appear to depend on the context of the behavioral evaluation (Mandairon et al., 2006).

### The role of VDCCs in generating beta oscillations

The persistence of beta oscillations after APV infusion implicates VDCCs in supporting beta oscillations, because AMPAR currents alone could probably not maintain sufficiently strong MC inhibition to support beta oscillations when NMDARs are blocked. In our model we found that beta power would stay relatively constant for moderate NMDA current suppression, but would increase for stronger NMDA current suppression (Fig. 1C, bottom). This is because the combination of NMDA and VDCC currents over-inhibited MCs, and a reduction of NMDA current shifted the system closer to a balance of excitation and inhibition, resulting in higher power oscillations. Though not significant, it is interesting to note that the LaOB showed some beta increases (Fig. 8A), which could potentially be attributed to a decrease in over-inhibition that occurs when NMDA is significantly blocked (assuming the system started in an over-inhibited state as shown in Fig. 1C).

An obvious next step would be to directly test the involvement of N-Type VDCCs in generating beta by infusing the N-type calcium channel blocker ω-conotoxin, but we leave this for future experiments. GCs are also known to express other VDCC subtypes, notably T-type channels which mediate Ca^2+^ spikes that can spread activity across the entire dendritic arbor to synchronize inhibition of all MCs connected to a given GC cell (Egger et al., 2005). The role that VDCCs play in OB oscillations is still being researched, but our results suggest that they are necessary for switching between gamma and beta oscillatory states in OB granule cells.

### Concluding Remarks

In summary, we found confirmation of our model’s main predictions, that reduced GC excitability can have a bidirectional effect on the power of OB beta oscillations, and that OB beta oscillations can be sustained independently of NMDARs. Going beyond the scope of our model, we also recorded PC beta oscillations. Though PC and OB beta oscillations had nearly identical frequency (Lowry and Kay, 2007; Kay and Beshel, 2010), we found that reductions in OB beta power through reduced GC excitability were not accompanied by significant reduction in PC beta power. Furthermore, OB-PC coherence was also not significantly reduced, even when OB beta power was dramatically reduced. It is possible that an OB-PC loop was sustaining PC beta, even though the beta component in the OB was very small under these conditions. The drug is most likely not targeting all GCs in the bulb equally, and some might be able to maintain an intact OB-PC loop. It is also possible that PC is just driving a small beta signal in the OB. Interestingly, a model of OB beta generation from a different group did not require OB and PC to coherently oscillate at beta frequency, but rather required a slow (theta frequency) modulation of GC excitability by PC inputs (David et al., 2015). Our model also did not involve an oscillating PC, and the persistent beta coherence we found in experiments suggests that the existence of OB-PC beta coherence is not itself sufficient for generating full-blown OB beta oscillations. While intact bidirectional OB-PC connections are required to generate beta oscillations, the GCs appear to control OB beta power, and to mediate the transition from gamma to beta oscillations.

GCs can exert at least four distinct types of inhibition onto MCs (Mouret et al., 2009). Dendrodendritic MC-GC connections can support local recurrent and lateral graded inhibition without GC somatic action potentials. But when GC action potentials are triggered, either by strong distal inputs or from MC or cortical inputs to the proximal dendrites near the soma, it is hypothesized that GCs switch to a global inhibition state, characterized by synchronous GABA release from all distal spines (Egger et al., 2005; Egger, 2008). The model developed by David et al. (2015) showed that switching between local and global inhibitory spiking GC states could drive transitions between gamma and beta oscillations, though the characteristics of GC firing patterns during beta oscillations have yet to be reported.

## Acknowledgments

We thank Selina Baeza-Loya and Cameron Westerback, for help in running some of the sessions and Vivian Nguyen for help with the high resolution figures.

